# Cationic antimicrobial copolymers reveal immunomodulatory properties in LPS stimulated macrophages *in vitro*

**DOI:** 10.1101/2025.04.08.647761

**Authors:** Sophie Laroque, James Harris, Santhosh Kalash Rajendrakumar, Vadim Vasilyev, Jaspreet Grewal, Robert Dallmann, Katherine E. S. Locock, Sébastien Perrier

## Abstract

Antimicrobial polymers, which have emerged as a promising alternative to antibiotics in the fight against antimicrobial resistance, are based on the design of cationic host defence peptides (CHDPs). Being a part of the mammalian innate immune system, CHDPs possess both antimicrobial and immunoregulatory effects to manage bacterial infections. However, the immunomodulatory effects of antimicrobial polymers remain largely unexplored. Within this work, a library of 15 copolymers were synthesised by reversible addition-fragmentation chain transfer (RAFT) polymerisation and their abilities to modulate pro-inflammatory pathways in LPS-activated murine and human macrophages were investigated. We found that two diblock copolymers with cationic units copolymerised with either apolar or hydrophilic comonomers appeared to have anti-inflammatory activity through suppression of the activation of the NF-κB signalling pathway, scavenging of reactive oxygen species and reduced production of the pro-inflammatory cytokine IL-6. Furthermore, the cationic-apolar copolymer exhibits significant antimicrobial activity against *P. aeruginosa*. Thus, this promising copolymer holds potential as a dual-action therapeutic, effectively combating bacterial infections while curbing prolonged inflammation and thereby preventing sepsis at the site of infection.

## Introduction

Antimicrobial resistance is one of the most significant challenges humanity is facing as we have entered the post-antibiotic era in which an increasing rate of multi-resistant bacteria strains coincide with a decreasing rate of development of new antibiotics to treat them.^1^ Over the last 30 years antimicrobial polymers have emerged as a potential alternative to antibiotics, due to their broad spectrum activity and mechanism of action.^2^ These cationic amphiphilic polymers are designed to mimic the ability of antimicrobial peptides (AMPs) to bind and penetrate the negatively charged bacterial cell surface, ultimately disrupting cellular structure.^3^

AMPs are a part of the innate immune system of multicellular organisms and play a central role in the defence against pathogenic microorganisms. These peptides have been studied extensively for their antimicrobial activities.^4–6^ However, further research over the past few decades has demonstrated immune-regulatory potential at sub-bactericidal concentrations, prompting a shift in their classification to ‘cationic host defence peptides’ (CHDPs).^7^ Therefore, these peptides primarily exhibit antimicrobial properties in physiological conditions through the modulation of the innate immune system, along with direct bacterial cell killing. Extensive research has highlighted the pivotal role of these peptides as immune modulators in the innate immune response through various mechanisms. Studies have shown that they are an essential modulator of the innate immune response, such as for example involvement in tissue and wound repair^8^, recruitment of immune cells to site of infection^9^, promoting bacterial clearance^10^, and reduce production of pro-inflammatory cytokines by removing endotoxins.^11^ CHDP’s have since then been investigated as both an antimicrobial agent^12^ and a potential treatment for chronic inflammation and sepsis.^13^

Sepsis is defined as uncontrolled harmful systemic inflammation which is induced by the prolonged presence of endotoxins after a bacterial infection. One such potent endotoxin is lipopolysaccharides (LPS), present on the outer membrane of gram-negative bacteria.^14^ LPS initiates a pro-inflammatory cytokine ‘storm’ by activating the toll-like receptor 4 (TLR4) and the nuclear factor kappa-light-chain enhancer of the activated B cells (NF-κB) pathway in the tissue-resident macrophages during bacterial infection, causing sepsis or septic shock^15 16^. The activation of the TLR4/NF-κB pathway leads to the secretion of proinflammatory cytokines, such as tumour-necrosis factor α (TNFα) and interleukin-6 (IL-6).^17^ This activation also results in the production of reactive oxygen species, such as nitric oxide (NO) and hydrogen peroxide (H_2_O_2_), which further activates the secretion of proinflammatory cytokines in the distal immune cells.^18^ These events lead to the activation and inflammation of other pro-inflammatory innate immune cells, including monocytes and neutrophils, which migrate to the site of infection. This cascade of events can cause multi-organ damage and, in severe cases, may lead to death^19^. CHDPs possess the ability to prevent sepsis by binding to the negatively charged moieties within LPS and blocking the initiation of the TLR4/NF-κB pathway,^11^ thus preventing the production of proinflammatory cytokines by directly interacting with macrophages.^20^

The effects of CHDPs are complex and can be both pro- and anti-inflammatory and their dysregulation can lead to immune dysfunction. For example, overexpression of the antimicrobial peptide LL-37, can lead to psoriasis.^21^ An absence of CHDPs can also contribute to disease, such as an increased rate of urinary tract infections.^22^ Therefore, should synthetic polymer mimics of these peptides be used in clinical settings, their potential immunomodulatory effects could be critical.

Interestingly, it has been shown that quaternized poly(isobutylene-*alt*-N-alkyl maleimide)s reduce the production of pro-inflammatory cytokines by binding to LPS and leading to a pseudo-aggregate formation.^23^ Only polymers with an increased hydrophobicity, however, were shown to bind to LPS and inhibit cytokine release in this study.^23^ Further systematic investigations are essential to more comprehensively assess the immunomodulatory effects of antimicrobial polymers while retaining their antimicrobial effects through for example varying their molecular weight as well as monomer composition.

Recently, we have demonstrated antimicrobial activity of copolymers synthesized via RAFT polymerisation against gram-negative and gram-positive bacteria.^3, 24–27^ Here, we address the immunomodulatory effect of these copolymers in innate immune cells such as macrophages. Specifically, we have investigated a library of 15 copolymers synthesised via RAFT polymerisation, based on the design of antimicrobial peptides. The monomer side chain functionalities were varied to obtain polymers with and without cationic units, as well as polymers with varying degrees of hydrophobicity. The molecular weight and the monomer distribution along the polymer chain were systematically varied to explore their potential impact on their biological properties. Their immunomodulatory activities were then investigated in LPS activated murine and human macrophages.

## Experimental Section

### Materials

4,4ʹ-Azobis(4-cyanovaleric acid) (ACVA), Acryloyl chloride, bis(*tert*-butoxycarbonyl)-2-methyl-2-thiopseudourea, Boc-anhydride (Fluka), chloroform (CHCl_3_), dimethyl sulfoxide-d_6_ (DMSO, 99.5 %), diethyl ether (≥ 99.9 %, inhibitor-free), dichloromethane (DCM), ethanol, ethyl acetate (EtOAc), ethylenediamine (99 %), hexane, magnesium sulfate (MgSO_4_), Müller-Hinton Broth type II (MHB cationic adjusted), *N*-isopropylacrylamide (NIPAM, 97 %), phosphate buffered saline (PBS) tablets, triethylamine (NEt_3_), trifluoro acetic acid (TFA), triton-X, 1,4-dioxane (≥ 99) were purchased from Sigma-Aldrich.

Corning Costar Flat Bottom Cell Culture Plates (Bottom: Flat, Clear, Lid: With Lid, Polystyrene, No. of Wells: 96, Sterile, Surface Treatment: Tissue-Culture Treated), Sodium Chloride and Suprasil® quartz cuvettes were purchased from Fisher Scientific.

Pre-wetted RC tubings 1 kD were purchased from Spectrumlabs. 2-((Butylthio)-carbonothioyl) thio propanoic acid (PABTC) and *N*-t-butoxycarbonyl-1,2-diaminoethane (BocAEAM) were synthesized and purified according to the reported literature.^27^ The bacterial isolates used (*Staphylococcus aureus* USA300 and *Pseudomonas aeruginosa* LESB058) were obtained from the library of Freya Harrison’s laboratory, School of Life Sciences, University of Warwick.

### Syntheses

#### Synthesis of (Propanoic acid)yl butyl trithiocarbonate (PABTC)

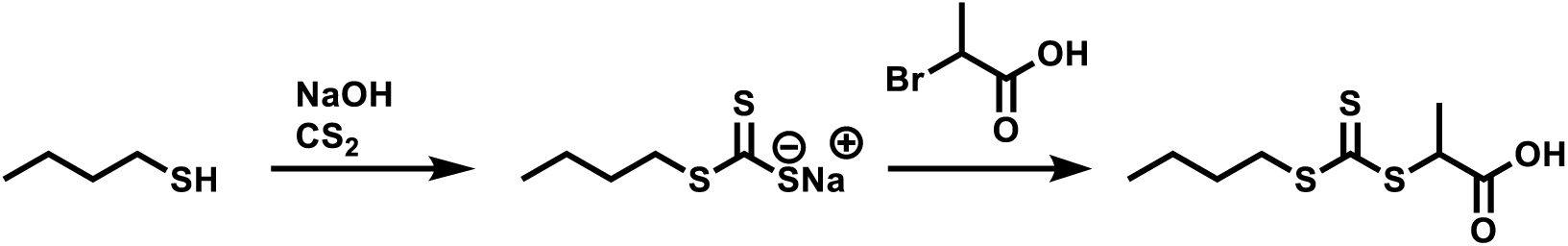

Sodium hydroxide (9.68 g, 0.242 mol, 1.1 eq) dissolved in 10 mL deionised water (50% w/w) was added dropwise to a solution of butanethiol (20 g, 0.22 mol, 1 eq) in acetone (11 mL) in a 500 mL round bottom flask, followed by addition of 40 mL deionised water and left to stir for 30 minutes at room temperature. Carbon disulfide (17.32 g, 0.228 mol, 1.025 eq.) was then added dropwise while stirring, and then left to stir for 30 minutes resulting in the solution turning into a yellow/orange colour. The mixture was then cooled with an ice bath to below 10 °C before adding 2-Bromopropionic acid (34.9 g, 0.228 mol, 1.025 eq) slowly and in a dropwise manner. followed by the slow addition of a 50% w/w aqueous NaOH solution (19.36 g, 0.484 mol, 1.00 eq) and left to stir overnight at room temperature. The solution is then diluted with a further 200 mL of deionised water and cooled to near 0 °C. 6M HCl was added dropwise until a pH 3 is reached which results in the precipitation of a yellow solid. The solid was filtered, and washed with water followed by recrystallisation in hexane yielding a yellow crystalline solid (yield 68%, 37.5 mmol).

^1^H NMR (400 MHz, CDCl_3_) δ = 4.87 (q, *J* = 7.4 Hz, 1H, C**H**(CH_3_)), 3.37 (t, *J* = 7.4 Hz, 2H, S-C**H_2_**-CH_2_**-**CH_3_), 1.74 – 1.65 (m, 2H, S-CH_2_-C**H_2_-**CH_3_), 1.63 (d, *J* = 7.4 Hz, 3H, CH(C**H_3_**)), 1.44 (d, *J* = 7.5 Hz, 2H, CH_2_-C**H_2_-**CH_3_), 0.94 (t, *J* = 7.3 Hz, 3H, CH_2_-CH_2_**-**C**H_3_**) ppm.

APT ^13^C-NMR (400 MHz, CDCl_3_) δ = 222.3 (**C**=S), 176.3 (**C**=O), 47.6 (**C**H(CH_3_)), 37.3 (S-**C**H_2_-CH_2_), 30.03 (S-CH_2_-**C**H_2_), 22.2 (CH_3_-**C**H_2_), 16.8 (CH(**C**H_3_)), 13.7 (**C**H_3_-CH_2_) ppm.

#### Synthesis of N-t-butoxycarbonyl-1,2-diaminoethane

A solution of ethylenediamine (26.71 mL, 400 mmol, 1 eq.) in 400 mL of DCM was added to a 1 L round bottom flask fitted with a pressure equalising dropping funnel. After the solution was cooled to 0 °C with an ice-bath, a solution of di-tert-butyl dicarbonate (8.73 g, 40 mmol, 0.1 eq.) in 200 mL DCM was added dropwise over two hours under stirring. The mixture was then allowed to warm to room temperature and left stirring overnight. The solvent was removed by rotary evaporation, and 100 mL of water was added to the residue. The white precipitate was removed by filtration, and the filtrate saturated with sodium chloride and extracted with ethyl acetate (3 x 60 mL). The combined organic phases were dried over sodium sulfate, filtered and then solvent was removed by rotary evaporation to yield a pale oil identified as *N*-t-butoxycarbonyl-1,2-diaminoethane (5.3 g, 32 mmol, 82% yield).

^1^H-NMR (400 MHz, CDCl_3_): δ = 5.1 (s, 1H, CH_2_-N**H**); 3.14 (m, 2H, C**H_2_**-NH), 2.78 (m, 2H, C**H_2_**-NH_2_), 1.37 (s, 9H,C-(C**H_3_**)_3_) ppm.

#### Synthesis of N-t-butoxycarbonyl-N’-acryloyl-1,2-diaminoethane

*N*-t-butoxycarbonyl-1,2-diaminoethane (5.3 g, 32 mmol, 1 eq.) and triethylamine (3.32 mL, 20 mmol, 0.7 eq.) were dissolved in 40 mL of chloroform in a 100 mL round bottom flask fitted with a pressure equalising dropping funnel and cooled to 0 °C with an ice-bath while stirring. Acryloyl chloride (3.05 mL, 40 mmol, 1.2 eq.) was dissolved with 60 mL chloroform and added dropwise over one and a half hours while stirring. After addition the mixture was allowed to warm to room temperature and left stirring for one hour. The solvent was removed under reduced pressure and dissolved in a minimum amount of chloroform. The solution was washed with 40 mL water, which was then extracted with chloroform (3 x 60 mL). After drying over sodium sulfate and filtration solvent was removed by rotary evaporation to yield a white powder identified as N-t-butoxycarbonyl-N’-acryloyl-1,2-diaminoethane (6.2 g, 28 mmol, 88% yield).

^1^H-NMR (400 MHz, CDCl_3_): δ = 6.28 (d, 1H, CH=C**H_2_**), 6.11 (q, 1H, C**H**=CH_2_), 5.66 (d, 1H, CH=C**H_2_**), 5.1 (s, 1H, CH_2_-N**H**); 3.46 (m, 2H, NH-C**H_2_**-CH_2_), 3.35 (m, 2H, NH-CH_2_-C**H_2_**), 1.46 (s, 9H, C-(C**H_3_**)_3_) ppm.

APT ^13^C-NMR (400 MHz, CDCl_3_): δ = 132 (**C**H=CH_2_) 127 (CH=**C**H_2_), 79 (O-**C**-(-CH_3_)_3_), 42 (-NH-**C**H_2_-CH_2_-), 40 (-NH-CH_2_-**C**H_2_-), 26 (O-C-(-**C**H_3_)_3_) ppm.

#### General procedure of RAFT polymerization of polyacrylamides

The monomers (NIPAm/DMA/BocAEAm), the PABTC chain transfer agent, 4,4ʹ-Azobis(4-cyanovaleric acid) (ACVA) and dioxane were added to a 7 mL glass vial equipped with a rubber septum and a magnetic stirrer bar to obtain a solution with a total concentration of 3 mol L^-1^. The solution was degassed with nitrogen for 15 minutes, and the reaction was heated in an oil bath to 85 °C. After 1.5 hours had passed the vial was removed from the oil bath and the reaction quenched by exposure to oxygen.

For the chain extensions the first block was redissolved in dioxane, and the 2^nd^ monomer and ACVA initiator added to make up a solution with a total concentration of 0.5 mol^-1^ mL.

#### Synthesis of poly(2-acrylamido-2-methyl-1-propanesulfonic acid) (pAMPs)

BDMAT (39 mg, 0.06 mmol), AMPS (622 mg, 3 mmol), phosphate buffer tablet solution (1.5 mL), sodium hydroxide (0.06 mmol, 2.4 mg) and VA-086 (2 x 10^-3^ mmol, 30 μL, 20 mg ml^-1^ stock prepared from phosphate buffer solution) were introduced to a 7 mL glass vial equipped with a rubber septum and a magnetic stirrer bar to obtain a solution with a final concentration of 1.5 mol L^-1^. The solution was degassed with nitrogen for 15 minutes, and the reaction was heated in an oil bath to 90 °C, for the duration of time required to reach nearly full conversion (2 hours). At the end of the reaction, the vial was removed from the oil bath and the reaction quenched by exposure to oxygen.

#### General Procedure for the Deprotection of Boc-Protected Polymers

The polymers were dissolved in 1.5 mL of dioxane and 1.5 mL of TFA, heated to 40 °C and left to stir for two hours. The TFA and dioxane was removed by precipitation in diethyl ether. Subsequently the polymers were dissolved in 10 mL of deionised water and dialysed against an aqueous solution of sodium chloride with three water changes every 3 hours, followed by a dialysis against water with water changes three times every three hours. Finally, the polymers were freeze-dried to remove water.

### Techniques

#### Size Exclusion Chromatography

An Agilent Infinity II MDS instrument equipped with differential refractive index (DRI), viscometry (VS), dual angle light scatter (LS) and variable wavelength UV detectors was used. The system was equipped with 2 x PLgel Mixed D columns (300 x 7.5 mm) and a PLgel 5 μm guard column. The eluent is DMF with 5 mmol NH_4_BF_4_ additive. Samples were run at 1 ml min^-1^ at 50 °C. Poly(methyl methacrylate) standards (Agilent EasyVials) were used for calibration between 955,000 – 550 g mol^-1^. Analyte samples were filtered through a nylon membrane with 0.22 μm pore size before injection. Respectively, experimental molar mass (*M_n_*, SEC) and dispersity (*Đ*) values of synthesized polymers were determined by conventional calibration and universal calibration using Agilent GPC/SEC software.

#### Nuclear magnetic resonance (NMR) spectroscopy

^1^H-NMR and ^13^C-NMR APT spectra were recorded on Bruker Avance 300 and 400 spectrometers (300 MHz, 400 MHz respectively). Deuterated chloroform, deuterated water and dimethyl sulfoxide-d_6_ were used as solvents for all measurements. Data analysis was performed using Mestrenova.

### *In Vitro* Analysis

#### Cell Culture

THP-1 monocytes (obtained from ATCC®) were cultured in RPMI 1640 media + 10% FBS, 2mM L-glutamine (GlutaMAX) and 1% Pen/Strep (complete media) and differentiated into macrophages in a 96 well plate with 5 ng/mL phorbol 12-myristate 13-acetate (PMA) for 48 hours at 37°C in 5% CO_2_. Mouse RAW264.7 and RAW264.7 cells stably transfected with the NF-kB responsive E-selectin promotor (ELAM, a generous gift from Professor Matthew Sweet, University of Queensland) that drives luciferase expression (under G418 selection) were cultured in in RPMI 1640 + 10% FBS, 2mM L-glutamine (GlutaMAX),1% Pen/Strep, 150 μg/ml G418 (geneticin) at 37°C in 5% CO_2_.

#### THP-1 macrophage treatment with LPS and copolymers

THP-1 cells were differentiated with PMA in a 96 well plate (10^5^ cells/well) as above, then treated with polymers dissolved in complete media at two 200 μg mL^-1^ and 50 μg mL^-1^ for 30 min, followed by stimulation with LPS (100 ng mL^-1^) for 6 h. After incubation cells were spun at 400 × *g* for 5 min and the supernatant transferred to a new 96 well plate and stored at -20 °C prior to analysis by ELISA and LDH Assays as described below. Treatment was repeated with a wider range for 4 copolymers (6.25 μg mL^-1^ – 200 μg mL^-1^) in duplicate in three independent experiments.

#### IL-6 Enzyme-linked Immunosorbent Assay (ELISA)

The concentration of IL-6 in THP-1 cell supernatants was measured with an ELISA kit (R&D Systems human IL-6 Duoset) according to manufacturer’s instructions. Plates were measured for absorbance at 450 nm, with a reference at 620 nm, on a ClarioSTAR microplate reader.

#### Cytotoxicity of polymers, using the LDH Assay for THP-1 cells

LDH concentration in THP-1 cell supernatants was determined with an LDH assay (Roche Cytotoxicity Detection Kit) according to manufacturer’s instructions. Absorbance was measured on a ClarioSTAR® microplate reader.

#### NF-kB luciferase assay in RAW 264.7 cells

In a 96 well plate, 10^5^ cells/well RAW 264.7-ELAM luciferase reporter cells in complete media were seeded and incubated for 1 hour at 37°C in 5% CO_2_. The cells were treated with copolymer solutions in RPMI media at different concentrations (6.25 μg mL^-1^ - 200 μg mL^-1^). The plate was left to incubate for 30 minutes, followed by stimulation with 10 ng mL^-1^ LPS for 4 hours at 37°C in 5% CO_2_. The cells were washed once with PBS and lysed by addition of 20 μL/well cell lysis buffer (Promega). After 5 minutes 100 μL/well Luciferase Assay Reagent (Promega) was added, and photoluminescence was read on the ClarioSTAR® microplate reader immediately.

#### Cell viability of RAW 264.7 cells after polymer treatment using the CellTiter 96® AQ_ueous_ One Solution Cell Proliferation Assay (MTS)

In a 96 well plate, 1x10^4^ cells/well of RAW 264.7 cells in complete DMEM media were seeded and incubated for 18 hrs at 37°C in 5% CO_2_. The cells were then treated with polymer samples in complete DMEM media at different weight concentrations (500 μg mL^-1^– 4 μg mL^-1^) and incubated for 24 hours at 37°C in 5% CO_2_. As controls cells were treated with DMEM media only. For assessing cell viability, 20 μL of MTS Reagent (CellTiter 96® AQueous One Solution Cell Proliferation Assay (Promega)) was added to each well and incubated for 4 hrs at 37°C in 5% CO_2_, and absorbance at 490 nm was measured in a Tecan Spark® 10M microplate reader.

#### ROS Assay with DCFH-DA dye

In a 96 well plate, 5x10^4^ cells/well of RAW 264.7 cells in complete DMEM media were seeded and incubated for 18 hrs at 37°C in 5% CO_2_. The cells were then treated with polymer samples in complete DMEM media at different weight concentrations (200 μg mL^-1^-– 12.5 μg mL^-1^) and incubated for 30 minutes at 37°C in 5% CO_2_, followed by stimulation with 1 μg mL^-1^ LPS in complete DMEM media and then left to incubate for 4 hours at 37°C in 5% CO_2_. As controls cells were treated only with DMEM media or LPS. After incubation cells were washed once with 1 mM HEPES buffer, followed by addition of DCFH-DA solution (25 µM) in 1 mM HEPES buffer and left to incubate for 30 minutes at 37°C in 5% CO_2_. Then the dye was removed, and cells were washed twice with 1 mM HEPES buffer and subsequently fluorescence intensities at 495 nm excitation and 529 nm emission as well as microscopy images of individual wells were imaged with a Cytation™ 3 plate reader and imager.

#### Minimum Inhibitory Concentration Assay

Minimum inhibitory concentrations (MICs) were determined according to the standard Clinical Laboratory Standards Institute (CLSI) broth microdilution method (M07-A9-2012).^28^ A single colony of bacteria in agar plates was chosen and dissolved in fresh cationic adjusted Mueller-Hinton broth (caMHB). The concentration of bacterial cells was adjusted by measuring the optical density at 600 nm (OD_600_) to obtain a 0.5 Mackfarland equivalent thus reaching a bacterial concentration of approximately 1 x 10^8^ colony forming unit per mL (CFU mL^-1^). The solution was further diluted by 100-fold to obtain a concentration of 1 x 10^6^ CFU mL^-1^. Polymers were dissolved in respective media and 50 μL of each polymer solution was added to micro-wells followed by the addition of the same volume of bacterial suspension, resulting in a final bacterial density of 5 x 10^5^ CFU mL^-1^. The micro-well plates were incubated at 37 °C for 18 hours. Then, the growth was evaluated by addition of 10 μL resazurin dye of each well leading to a final concentration of 0.5 mg mL^-1^ The plates were incubated for 30 minutes at 37 °C and a noticeable change of colour could be observed where bacteria cells grew (pink colour) while the suspension remains blue in wells with non-detectable growth. Resazurin was prepared at 0.5 mg mL^-1^ stock in PBS and the solution was filter sterilised (0.22 µm filter). The solution was stored at 4 °C and covered in foil for a maximum of 2 weeks after preparation. The protocol was followed as described before by M. Elshikh *et al*.^29^

#### Fluorescent Confocal Microscopy

RAW 264.7 cells were seeded in an 8 well chamber slide in complete DMEM media at a concentration of 5x104 cells/well and left to incubate overnight in 37°C at 5% CO_2_ to let cells adhere. Media was removed, followed by addition of 250 μL copolymers stock solutions in DMEM media at weight concentration of 50 μg mL^-1^ and left to incubate for 30 minutes, followed by addition of 250 μL of 2 μg mL^-1^ LPS solution (in DMEM) and then left to incubate for 4 hours at 37°C in 5% CO_2_. After incubation media was removed and cells gently washed with PBS twice. 100 μL of 4% paraformaldehyde was added and left to incubate for 15 minutes at room temperature. The paraformaldehyde was removed, followed by removing the top wells to yield the glass slide, which was left to dry at room temperature. One drop of DAPI containing antifade solution was added per well, and the cover slide placed on top. Finally, the sample was left to dry overnight, and imaging was performed on a Zeiss LSM 880 fluorescence microscope. For the kinetic experiment no LPS was added and copolymer solution at weight concentrations of 50 μg mL^-1^ was added at several time points.

#### Transcriptomic analysis

For the RNA sequencing experiment, RAW247.9 cells were plated in a 96 well plate and treated with polymer at a concentration of 50 μg mL^-1^ for 30 minutes, followed by stimulation with 1 µg/mL LPS and left to incubate for 4 hours 37°C in 5% CO_2_. Then, cells were washed, trypsinized and collected to perform the total RNA extraction with the RNAeasy plus kit (Quiagen) and genomic DNA removal columns following the manufacturer’s instructions. Libraries were prepared after poly(A) enrichment and then sequenced on an Illumina NovaSeq using pair-ended 150bp kit to a depth of at least 20 mi reads at Azenta (Germany). Gene expression tables were generated from FASTQ data output by aligned to the mouse genome GRCm39 using Kallisto on the BioJupies platform.^30^ Gene differential expression (DE) analysis was conducted using the standard DESeq2 pipeline.^31^ Variance stabilising transformation was used for outlier detection and PCA / heatmap visualisations. In addition, values for heatmap generation were normalised by DESeq2’s median of ratios value and then z-transformed and Euclidean distance with complete linkage was used for clustering. DE genes were identified using FDR cutoff of 0.05 and absolute log2-fold change cutoff of 1.5. For the downstream gene set enrichment analysis, empirical bayes shrinkage for fold-change estimation was used.^32^

Gene Set Enrichment Analysis of KEGG pathways and visualisations were done using clusterProfiler package (gseKEGG) with a p value cutoff of 0.01.^33^

Transcription factor activity estimation was conducted using the decoupleR package. Briefly, CollecTRI containing a curated collection of TFs, their transcriptional targets, and the associated mode of regulation weights, was used as a prior knowledge database. A univariate linear model (ulm) was run for each sample and each TF in the network with the estimated t-value of the slope being the assigned score.^34^ For the LPS-Polymer vs LPS comparison we used DESeq2’s test statistic values of the DE genes between these two conditions to fit ulm.

## Results and Discussion

### Synthesis of Copolymer Library by RAFT Polymerisation

To assess how different side-chain functionalities might affect the immunomodulatory effects, a copolymer library was synthesised by reversible addition-fragmentation chain-transfer polymerisation (RAFT). A library of 15 polymers were synthesised (**Figure 1**), using a total of 5 different acrylamide monomers to investigate the effects of the balance of hydrophobicity and hydrophilicity and the influence of cationic and anionic charges. We chose acrylamides as the monomer class, as they enable fast synthesis of well-defined structures and result in highly stable polymers which are resistant to hydrolysis and degradation by enzymes.^35^ PABTC was chosen as the chain transfer agent (CTA) as trithiocarbonate CTAs have shown good results for the polymerisation of acrylamides and appear hydrolytically stable.^36^

**Figure 1:**
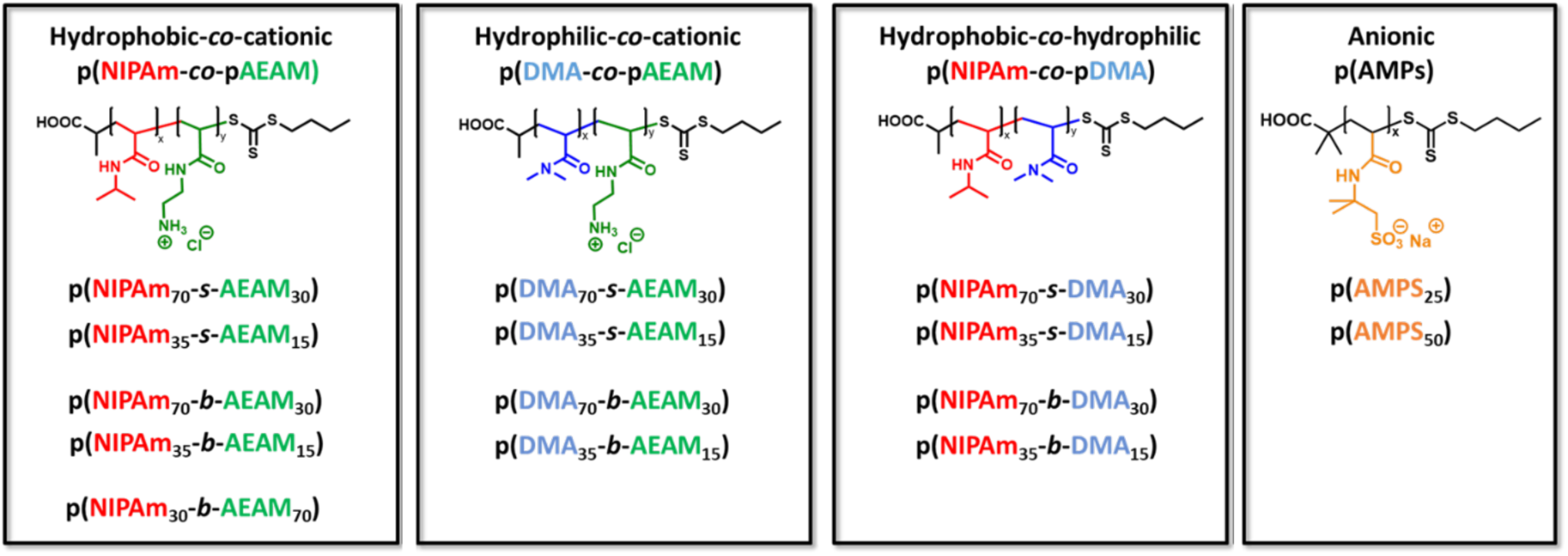
Copolymer Library to be synthesized by RAFT Polymerisation and investigated for their immunomodulatory activities. The properties are systematically changed to investigate influences of hydrophobicity, hydrophilicity, and cationic/anionic charges.

Within our group we have conducted several studies on antimicrobial properties of polyacrylamides,^3, 24, 26, 27^ comprised of *N-*isopropyl acrylamide (NIPAm) and aminoethyl acrylamide (AEAm). AEAm was chosen as a lysine mimic and NIPAm as a leucine mimic, amino acids commonly found in antimicrobial peptides,^12^ and it has resulted in copolymers with high antimicrobial activity and good hemocompatibility. NIPAm is a water soluble monomer, however it has an apolar side-chain and confers an overall amphiphilic character to the final structure when copolymerised with a cationic monomer. Therefore, within the context of this study it is referred to as ‘hydrophobic’. pNIPAm is known to have a lower critical solution temperature at 32 °C,^37^ however it has been shown in previous studies^3, 24, 37^ that through copolymerisation with a hydrophilic or cationic comonomer the LCST is raised far above the physiological temperature of 37 °C at which these copolymers are tested for their biological activity. Therefore, no interference due to aggregation is expected when determining their anti-inflammatory and antimicrobial properties at this temperature.

In order to test if these antimicrobial polymers have immunomodulatory properties, we synthesised hydrophobic-cationic copolymers based on NIPAm and AEAm (**Figure 1**). For one of these polymers the ratio of cationic units was changed, from 30% to 70%, to determine how increasing cationic units affect immunomodulatory properties. To establish how exchanging NIPAm for a more hydrophilic monomer will impact the structure-activity relationship, NIPAm was replaced by dimethyl acrylamide (DMA) to generate hydrophilic-cationic polymers. Finally, to probe if the cationic charges are essential, a third combination of monomers saw the AEAm units replaced by DMA, thus resulting in hydrophobic (pNIPAM) – hydrophilic (pDMA) polymers. In addition, homopolymers with the negatively charged monomer 2-acrylamido-2-methylpropane sulfonic acid (AMPs) were synthesised, as pAMPs is known to mimic heparin,^38^ which can have anti-inflammatory effects.^39^ For each of these polymer categories the molecular weight was varied by targeting a degree of polymerisation (DP) of 50 and 100. For the pAMPs the DP targeted were 25 and 50, which approximately matches the molecular weight for the copolymers with DP 50 and 100. Furthermore, the copolymers structures were changed from statistical copolymers to diblock copolymers, as it has been shown to have a great impact on interactions of polymers with cell membranes.^27, 40^ To summarise, the polymer library comprises 15 polymers which are grouped into 4 categories: hydrophobic-cationic copolymers are numbered as copolymers 1-5, hydrophilic-cationic as copolymers 6-9, hydrophobic-hydrophilic as copolymers 10-13 and anionic homopolymers as 14 and 15 (**Table 1**).

**Table 1:** SEC (DMF SEC, PMMA Standard) and ^1^H-NMR (300 MHz, CDCl_3_) results for linear copolymers.

| | Polymer | DP | Conversion [%] | $M_{wNMR}$<br>[g mol<sup>-1</sup>] | $M_{nSEC}$<br>[g mol<sup>-1</sup>] | $\bar{D}$ |
| --- | --- | --- | --- | --- | --- | --- |
| 1 | p(NIPAm<sub>35</sub>-s-BocAEAm<sub>15</sub>) | 50 | 99 | 7400 | 9200 | 1.12 |
| 2 | p(NIPAm<sub>70</sub>-s-BocAEAm<sub>30</sub>) | 100 | 99 | 14600 | 14800 | 1.14 |
| 3 | p(NIPAm<sub>35</sub>-b-BocAEAm<sub>15</sub>) | 50 | 99 | 7400 | 9700 | 1.18 |
| 4 | p(NIPAm<sub>70</sub>-b-BocAEAm<sub>30</sub>) | 100 | 99 | 14600 | 13000 | 1.10 |
| 5 | p(NIPAm<sub>30</sub>-b-BocAEAm<sub>70</sub>) | 100 | 99 | 18600 | 18400 | 1.23 |
| 6 | p(DMA<sub>35</sub>-s-BocAEAm<sub>15</sub>) | 50 | 99 | 6900 | 11400 | 1.13 |
| 7 | p(DMA<sub>70</sub>-s-BocAEAm<sub>30</sub>) | 100 | 99 | 13600 | 20000 | 1.16 |
| 8 | p(DMA<sub>35</sub>-b-BocAEAm<sub>15</sub>) | 50 | 99 | 6900 | 10100 | 1.20 |
| 9 | p(DMA<sub>70</sub>-b-BocAEAm<sub>30</sub>) | 100 | 99 | 13600 | 16900 | 1.16 |
| 10 | p(NIPAm<sub>35</sub>-s-DMA<sub>15</sub>) | 50 | 99 | 5700 | 8700 | 1.10 |
| 11 | p(NIPAm<sub>70</sub>-s-DMA<sub>30</sub>) | 100 | 99 | 11100 | 16000 | 1.13 |
| 12 | p(NIPAm<sub>35</sub>-b-DMA<sub>15</sub>) | 50 | 99 | 5700 | 8900 | 1.13 |
| 13 | p(NIPAm<sub>70</sub>-b-DMA<sub>30</sub>) | 100 | 99 | 11100 | 14900 | 1.18 |
| 14 | P(AMPS)<sub>25</sub> | 25 | 99 | 5400 | n.a. | n.a. |
| 15 | P(AMPS)<sub>50</sub> | 50 | 99 | 10600 | n.a. | n.a. |

All 15 copolymers were successfully synthesised to quantitative monomer conversion with consistently low dispersities, below 1.3 (**Table 1**, **Figure 2**). During synthesis of the cationic copolymers, the AEAm monomer was protected with a Boc group to prevent aminolysis which was removed post-polymerisation by treatment with trifluoroacetic acid (TFA) followed by dialysis against aqueous sodium chloride solution to exchange the counterion.

**Figure 2:**
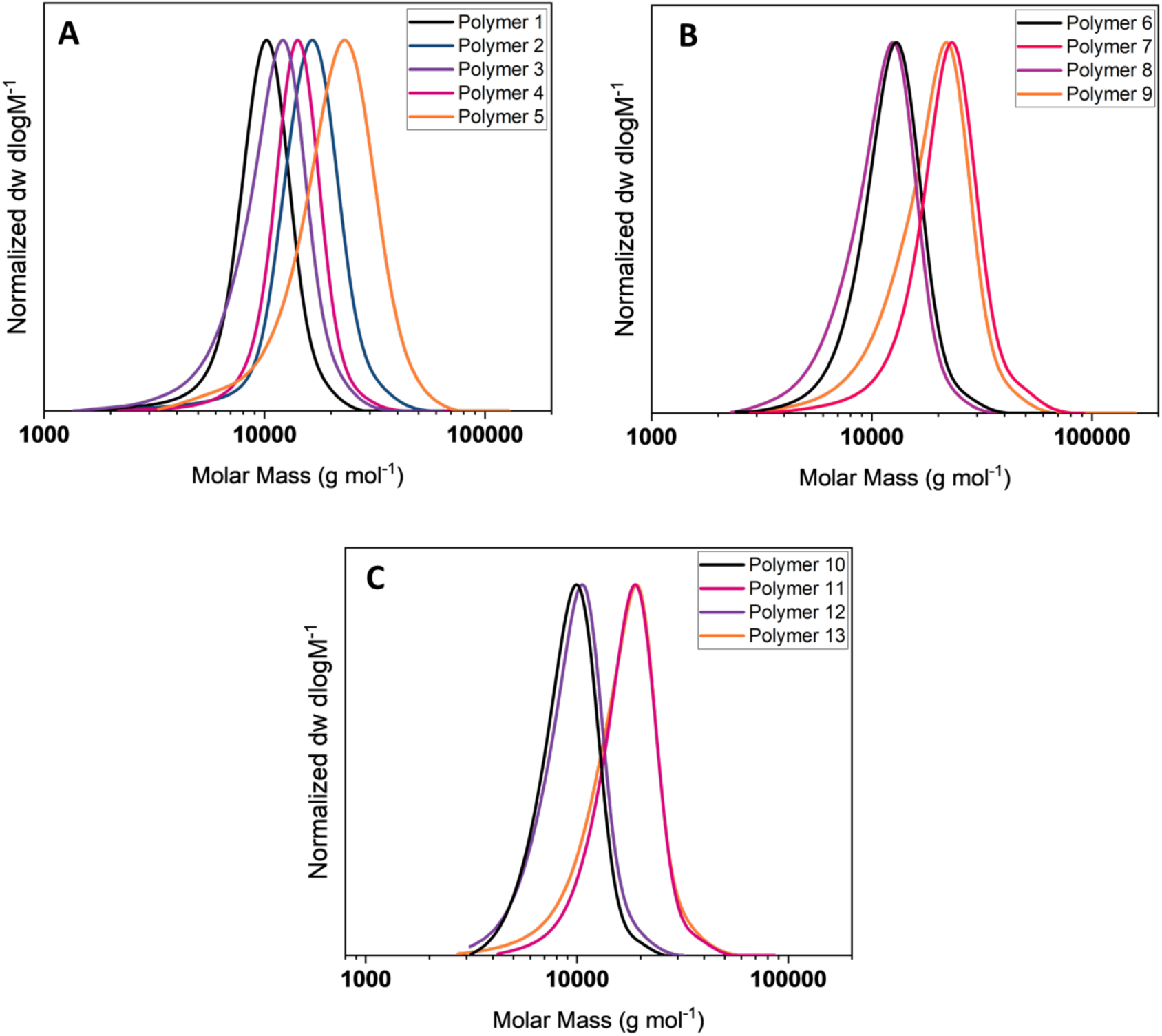
SEC Traces of Linear Copolymers (DMF GPC, PMMA Standard), **(A)** – Polymers 1-5, **(B)** Polymers 6-9, **(C)** – Polymers 10-13.

### Screening of copolymer library for immunomodulatory activities and cytotoxicity

THP-1 monocytes,^41^ a human monocytic cell, line are widely used as models to investigate the anti-inflammatory responses of compounds *in vitro*.^41^ Upon treatment with LPS, they demonstrate an inflammatory phenotype, with increased expression of a wide range of inflammatory genes and production and release pro-inflammatory cytokines, such as IL-6.^42^

The effects of polymers on the secretion of IL-6 after LPS stimulation in THP-1 derived macrophages were screened using enzyme-linked immunosorbent assay (ELISA). A Lactate dehydrogenase (LDH) assay was performed simultaneously in order to confirm that any reduction in secretion of IL-6 was not due to cytotoxic effects. The IL-6 levels in the polymer-treated cells were compared to untreated and LPS-treated cells, serving as positive and negative controls respectively. LDH release by polymer treated cells was compared to LPS-treated cells, and deemed as cytotoxic if the change in LDH release was far higher in direct comparison (10% LDH release for LPS).

The copolymers 10-15 (hydrophobic-hydrophilic and anionic copolymers) showed no obvious effects on IL-6 release and likewise no discernible effect on LDH release compared to LPS alone, suggesting no cytotoxicity (Figure 3).

**Figure 3:**
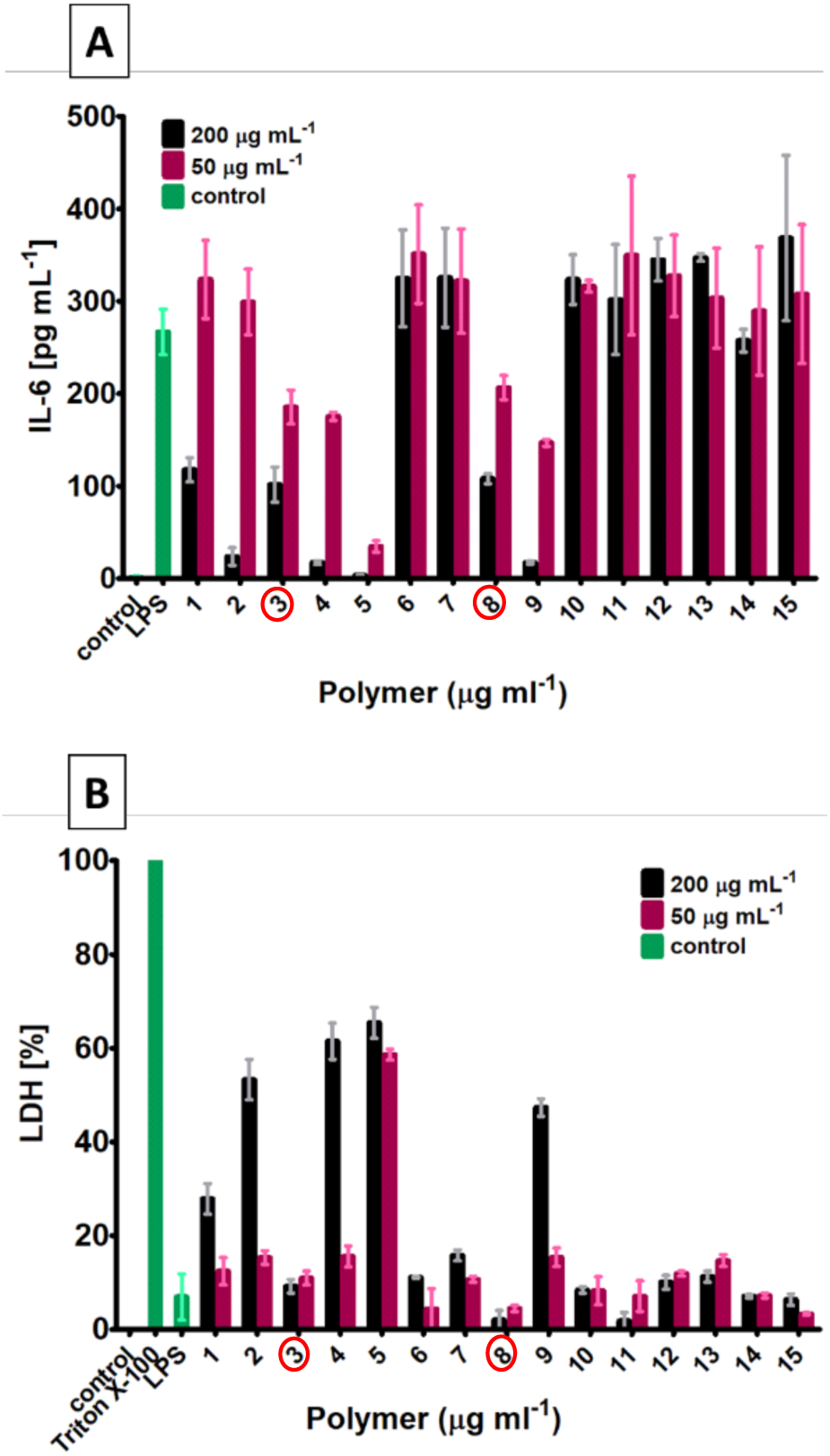
Effect of polymers on the secretion of IL-6 after LPS stimulation (**3A**), and cytotoxic effects measured by determination of LDH release in in THP-1 derived macrophages (**3B**). Two concentrations were measured for each copolymer, and each sample was measured in duplicates. Control sample are untreated cells. Copolymers marked red were deemed non-cytotoxic and reduced IL-6 in comparison to LPS control.

For the copolymers 6-9 (hydrophilic-cationic library), copolymers 8 and 9 had inhibitory effects on LPS-induced IL-6 release (Figure 3A). Copolymer 8 had no observable effect on LDH release, indicating that cytotoxicity was not induced (Figure 3B). However, copolymer 9 did increase LDH release (Figure 3B), suggesting that its effects on IL-6 release might be due to cytotoxicity. Copolymers 6 and 7, both with statistical monomer distribution had no obvious effects on LDH or IL-6 release.

For copolymers 1-5, copolymer 5, which contains 70% cationic units, potently reduced IL-6 release at both 50 and 200 μg ml^-1^, but also exhibited the highest levels of cytotoxicity within this library at both concentrations (Figure 3B). Copolymer 4 was likewise highly cytotoxic at the highest dose (200 μg ml^-1^), but not at 50 μg ml^-1^ (Figure 3B), a dose at which it showed some inhibitory effect on IL-6 release (Figure 3A). The statistical copolymers 1 and 2 both had inhibitory effects on IL-6 release at 200 μg ml^-1^ (Figure 3A) but were both cytotoxic at this dose (Figure 3B). The smaller diblock copolymer 3, on the other hand, exhibited inhibitory effects on LPS-induced IL-6 release without any effect on LDH release.

This initial screening assay would suggest that the main factor contributing to cytotoxicity is the cationic charge. A second factor leading to an increased cytotoxicity is hydrophobicity, when comparing the two cationic libraries, as replacing NIPAm with DMA greatly reduced amount of LDH release. Finally, an increased molecular weight also correlated with increased cytotoxicity, as did a diblock rather than a statistical structure. These findings are in agreement with previous studies on the toxicity of these polyacrylamides against mammalian cell lines.^27^

Regarding the IL-6 release a clear pattern emerged, suggesting that cationic charges are necessary to inhibit the release of IL-6, as the non-cationic polymers showed no activity. Furthermore, diblock copolymers reduced IL-6 secretion further then statistical copolymers which lead to the identification of the two lead compounds polymer 3 and polymer 8.

These results indicate that the anti-inflammatory properties are tied to the cationic charges and are therefore important when considering the mode of action. We hypothesise that the cationic charges may enable the polymers to bind to negative moieties of LPS as well as bind and insert into the macrophage cells thereby disrupting pro-inflammatory pathways such as NF-kB and reducing IL-6 release.

To conclude diblock copolymers 3 p(NIPAm_35_-b-BocAEAm_15_) and 8 p(DMA_35_-b-BocAEAm_15_) maintained the balance between a significant reduction of IL-6 release in combination with a low cytotoxicity at both concentrations and were further investigated for their potential as an anti-inflammatory copolymer.

### Investigation of immunomodulatory effects of promising copolymers 3 and 8

To confirm the anti-inflammatory effects of copolymers 3 and 8 observed in the initial screening, and further refine a potential dose-activity relationship, we tested their effects on LPS-induced IL-6 release by THP-1 cells at concentrations between 12.5 – 200 μg ml^-1^, along with copolymer 12 and 14 as inactive controls (**Figure 4**). The results confirmed the significant suppressive effects of both copolymers 3 and 8 on LPS-induced IL-6 release.

**Figure 4:**
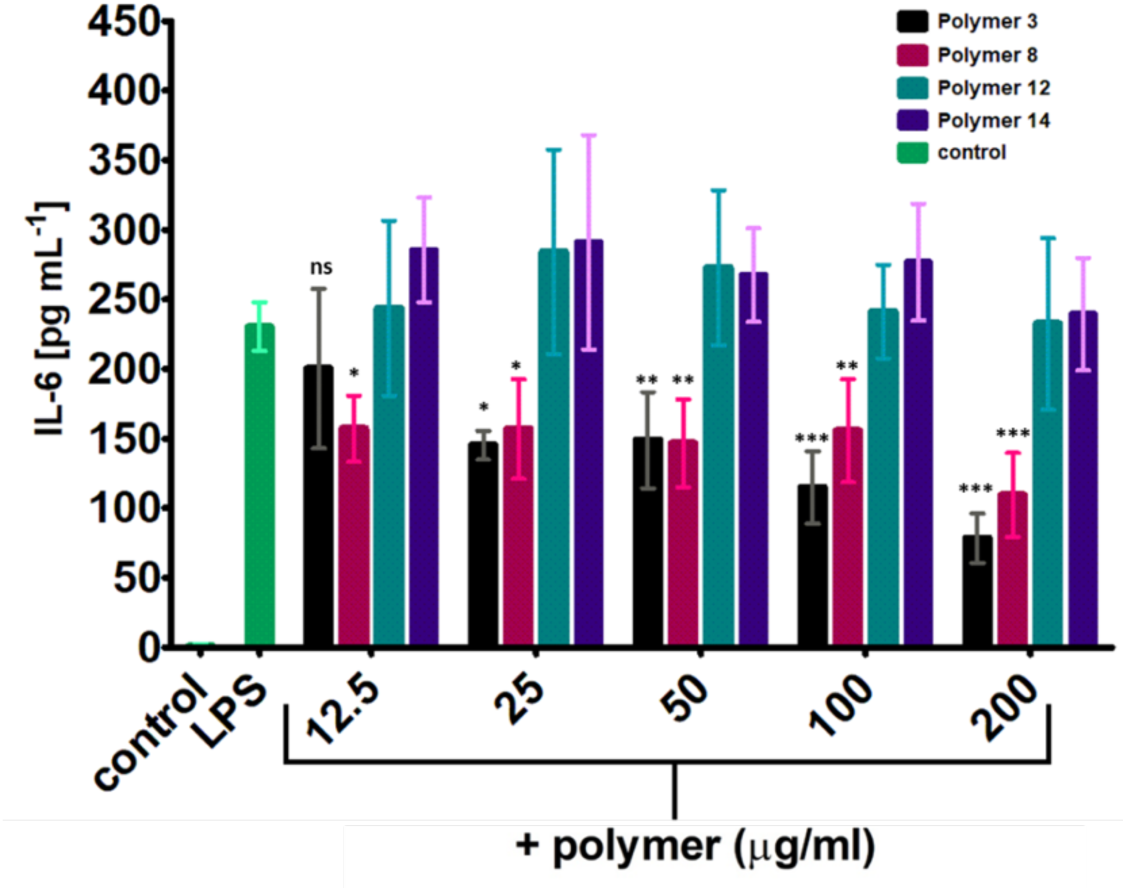
Effects of promising copolymers and control polymer on LPS stimulated THP-1 macrophages on pro-inflammatory cytokine release (IL-6) with a wider concentration range (200 μg mL^-1^ – 12.5 μg mL^-1^). Each copolymer was measured three times in triplicate. Analysed by two-way ANOVA, n=6 per group, Bonferroni post hoc test compared with LPS control: p<0.001 (***) for both Polymer 3 and 8, p < 0.05 (n.s.) for Polymer 12 and 14.

Next, the ability to suppress the activation of the NF-κB pathway in LPS activated NF-κB reporter murine (Luc)-RAW 264.7 cells was assessed.

A significant reduction of NF-κB activity was observed for both copolymer 3 and 8 at concentrations between 6.25 – 200 μg mL^-1^ (Figure 5A). At the highest measured concentration for copolymer 3 (200 μg mL^-1^) the activity of NF-κB was reduced by 83%, compared to LPS alone. In order to confirm that copolymers 3 and 8 are not cytotoxic and potentially reducing the activation of NF-κB by killing the cells, an MTS assay was performed, which clearly demonstrated no effect of the copolymers on cell viability and number (Figure 5B) within the concentration range of 2-500 μg mL^-1^. These results demonstrate the anti-inflammatory properties are consistently observed in both murine and human macrophage cell lines with two different assays.

**Figure 5:**
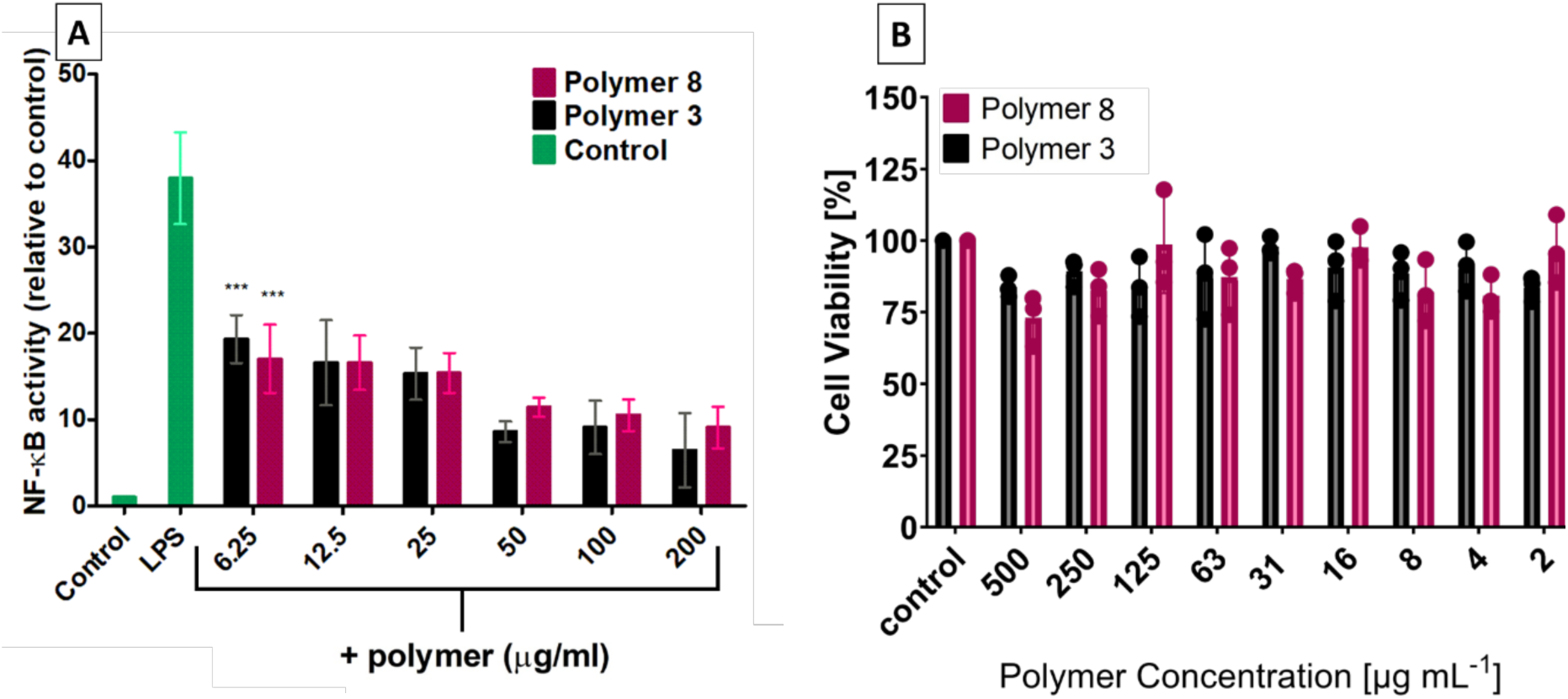
**A:** Effects of copolymers 3 and 8 on LPS stimulated Mouse RAW264.7 cells (stably transfected with the NF-κB responsive E-selectin promotor (ELAM)) on NF-κB activation with a wider concentration range (200 – 12.5 μg mL^-1^). Analysed by two-way ANOVA, n=6 per group, Bonferroni post hoc test compared with LPS control: p<0.001 (***) for both copolymer 3 and 8 at all concentrations measured. All concentrations were significant (***). (**B:** MTS Assay to determine cytotoxicity of copolymer 3 and 8 against RAW 274.7 cells with a wider concentration range (500 - 2 μg mL^-1^).

Investigation of ROS levels in RAW 264.7 cells after treatment with copolymers 3 and 8

The anti-inflammatory response generated by the polymers may affect intracellular reactive oxygen species (ROS) levels. Therefore, ROS levels in LPS stimulated RAW 264.7 cells post-polymer treatment were detected using 2ʹ,7ʹ,-dichlorofluorescein diacetate (DCFH-DA) fluorescence.

For the copolymer treated samples ROS levels were significantly reduced, by over 50% compared to the LPS control (Figure 6B). Slightly higher ROS levels were observed for copolymer 8 compared to copolymer 3. Microscopy images of the macrophages (Figure 6A) show that the fluorescence is emitted from the cells only, and confirms a higher intensity of fluorescence for the LPS treated cells compared to untreated and LPS plus copolymer treated samples. These results demonstrate that these copolymers can reduce ROS levels in LPS-stimulated macrophages, which further confirms their anti-inflammatory properties. When looking at hydrophobic versus hydrophilic no significant difference could be observed regarding the effects on IL-6 release, NF-κB activation and ROS reduction.

**Figure 6:**
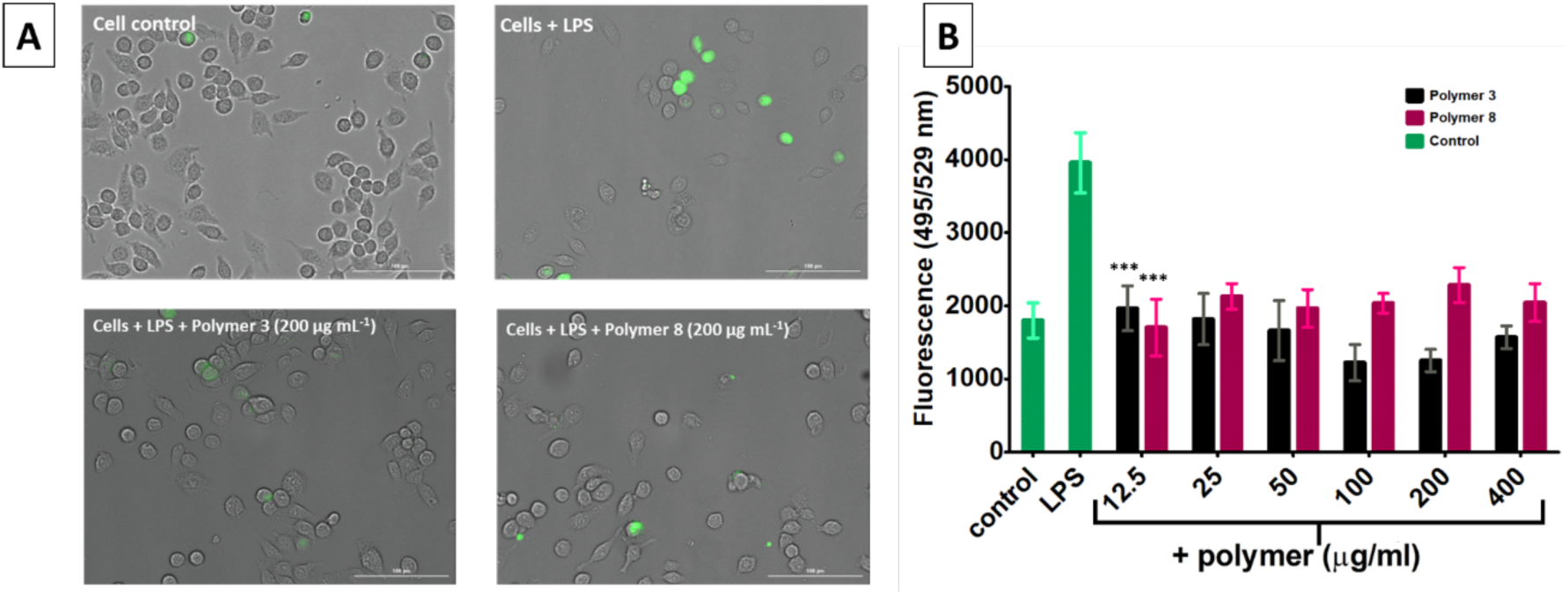
Effects of promising copolymers on production of intracellular reactive oxygen species (ROS) using the oxygen free radical acceptor 2ʹ,7ʹ,-dichlorofluorescein diacetate (DCFH-DA) in LPS stimulated Mouse RAW 264.7 macrophages. **A:** Microscopy Images were obtained, green fluorescence is indicative of ROS formation. **B:** Fluorescence that is representative of intracellular ROS level was measured at 495 nm excitation and 529 emission in a Cytation 3 Cell Imaging Plate Reader. 2-way ANOVA p<0.001 for both Polymer 3 and 8 (***) at all concentrations. Bonferroni post hoc test compared with LPS control.

Cell-uptake of copolymers 3 and 8 in LPS-activated 264.7 macrophages

To verify cellular uptake in macrophages, both copolymers 3 and 8 were functionalised with Cy-5, a far-red fluorophore to analyse the cellular uptake in DAPI stained RAW 264.7 cells with confocal microscopy (Figure 7). The cells were incubated with the Cy-5 copolymers for 30 minutes, then treated with LPS and left to incubate for 4 hours to replicate the conditions used in the previous assays.

**Figure 7:**
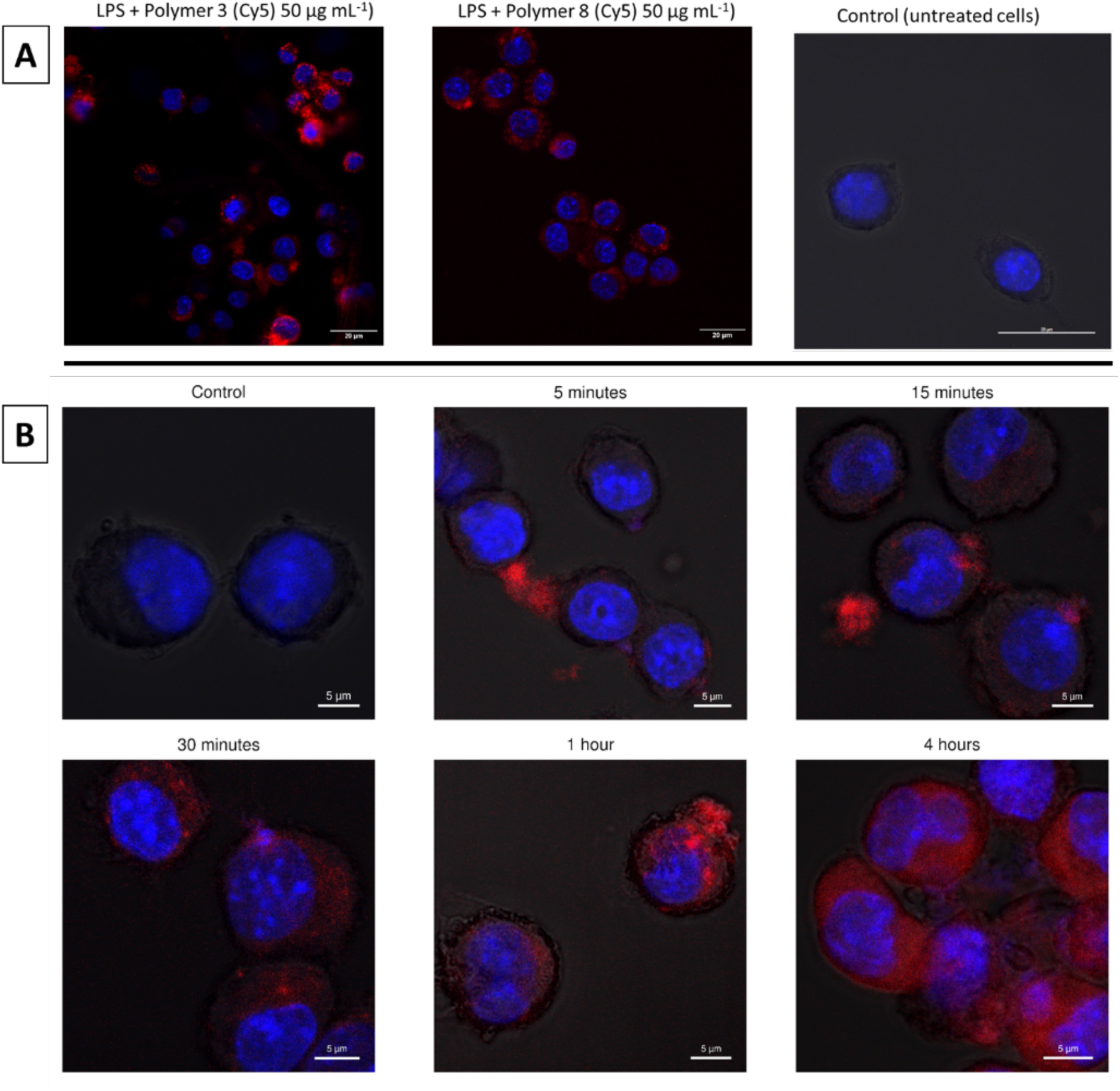
**A**: DAPI stained RAW 264.7 macrophages treated with Cy5 labelled copolymers for 4 hours at 50 μg mL^-1^. **B:** DAPI stained RAW 264.7 macrophages treated with Cy5 labelled copolymer 3 at 50 μg mL^-1^ at different time points. Control sample treated with DAPI only.

The images obtained clearly indicate that the copolymers are taken up by the RAW 264.7 macrophages. By examining the Z plane of these images (Supplementary Figure S8 and S9) it is confirmed that the copolymers have entered the cells and are not just aggregated on the surface. To further investigate cell-uptake the kinetics of cell uptake were identified for the lead compound copolymer 3 using a time course series. After 5 and 15 minutes some copolymer seems to have aggregated to the cell surface, however not much can be observed within the cell. After approximately 30 minutes most of the copolymer appears to be taken up by the cells shown in this sample. After 1 – 4 hours the concentration of copolymer increases further.

This experiment demonstrates that after the 30 minutes incubation prior to stimulation with LPS a significant amount of copolymer 3 is taken up by the cells. This is relevant when discussing the possibilities of interaction between LPS and the copolymer prior to activation of the pathways. As copolymer 3 is based on the design of CHDPs, we can look at their mode of action to draw potential conclusions on the anti-inflammatory properties observed in this study. CHDPs are known for acting as an anti-endotoxin by binding to the negatively charged LPS and preventing it from binding for example to TLR4 receptors on macrophages.^43^ This could therefore be the main mode of action of our copolymers in this study, given that only cationic copolymers showed anti-inflammatory properties.

However since the copolymers are readily taken up by the cells within 30 minutes incubation (Figure 7B), some intracellular effects cannot be excluded. Furthermore CHDPs are known to have multimodal functions, therefore the same could be the case for our copolymers. As an example, the CHDP LL-37 has been shown to both bind to LPS acting as an anti-endotoxin, as well as reducing the translocation of the NF-κB subunits p50 and p65 to the nucleus by over 50%.^43^ Further work is planned to investigate if intracellular effects are taking place.

The final step was to compare the antimicrobial activity of these two copolymers. Previous work in our group^3, 24, 25, 27^ has established significant antimicrobial activity for p(NIPAm-*b*-AEAm) copolymers. Therefore the assessment of biological activity towards bacteria was focussed on the two copolymers which demonstrated anti-inflammatory activity combined with low cytotoxicity within the scope of this work.

### Antimicrobial activity of copolymers 3 and 8

In order to evaluate antimicrobial activity of the two promising copolymers the minimum inhibitory concentration (MIC) was determined (using the cell viability dye resazurin).^29^ Two bacterial species were chosen for this study, a gram negative *P. aeruginosa strain* (LESB58), a clinical isolate found in chronic CF lung infections^44^, and a gram-positive *S. aureus* strain (USA 300), a methicillin-resistant strain and strong biofilm former (Table 2).

**Table 2:** Antimicrobial activity of copolymers 3 (p(NIPAm-b-AEAm)_50_) and copolymer 8 (p(DMA-b-AEAm)_50_) against *S. aureus* (USA 300) and *P. aeruginosa* (LESB58).^a^.

| Polymer | MIC (*S. Aureus*)<br>[$\mu\text{g mL}^{-1}$] | MIC (*P. Aeruginosa*)<br>[$\mu\text{g mL}^{-1}$] |
| --- | --- | --- |
| Polymer 3 | 256 | 32 |
| Polymer 8 | 512 | >512 |

<sup>a</sup> MICs values expressed in $\mu\text{g mL}^{-1}$ of the copolymers tested in caMHB against *S. aureus* USA300 and *P. aeruginosa* LESB58. Three independent biological experiments were performed on different days, and the highest MIC value was reported. Copolymers were tested in the concentration range of $8 \mu\text{g mL}^{-1}$ – $512 \mu\text{g mL}^{-1}$ .

For the copolymer 3 (p(NIPAm-b-AEAm)_50_), antimicrobial activity can be observed against both strains whereas copolymer 8 showed no activity against *P. aeruginosa* and exhibited an MIC only at the highest concentration measured against *S. aureus*. The overall lowest MIC was observed for copolymer 3 against *P. aeruginosa* with a value of 32 μg mL^−1^. When comparing the antimicrobial activity of the two promising copolymers the importance of the NIPAm block to confer antimicrobial activity becomes clear, as switching to DMA clearly reduced activity towards both strains. This is probably due to the increased hydrophobicity of pNIPAm compared to pDMA, allowing better penetration of the bacterial membrane.

Furthermore, copolymer 3 exhibits an MIC of 32 μg mL^-1^ against *P. aeruginosa* which is well above the concentration of copolymer at which anti-inflammatory effects were observed (6.25 μg mL^-1^). This is in agreement with observations that cationic host defence peptides have immunomodulatory effects at sub-MIC concentrations.^45^ This shows that these copolymers appear to be effectively mimicking CHDPs. Copolymer 3 is therefore identified as the lead compound with both antimicrobial and anti-inflammatory properties combined with low cytotoxicity.

Based on the significant activity of copolymer 3 against *P. aeruginosa* it appears to be the most promising compound within this study, as it exhibits both high antimicrobial activity and significant anti-inflammatory properties. It could therefore potentially be used as a topical treatment in wound infections for example, thereby both killing bacterial cells and acting as an anti-endotoxin, preventing prolonged inflammation and sepsis.

Differential gene expression induced by copolymer 8 in LPS activated RAW264.7 macrophages.

Next, we assessed the gene expression profile of copolymer 3-treated, LPS-activated RAW 264.7 cell line, given it demonstrated dual anti-inflammatory and antimicrobial activity. Principal component analysis (PCA) distinctly segregated different treatment groups, apart from the control and polymer-only treatment, which clustered together (Figure 8A). DESeq2 analysis revealed 100 uniquely down-regulated and 175 up-regulated genes in response to LPS, whereas in the presence of both LPS and polymer treatment, only 60 genes were down-regulated and 32 up-regulated (Figure 8B) compared to control.

**Figure 8:**
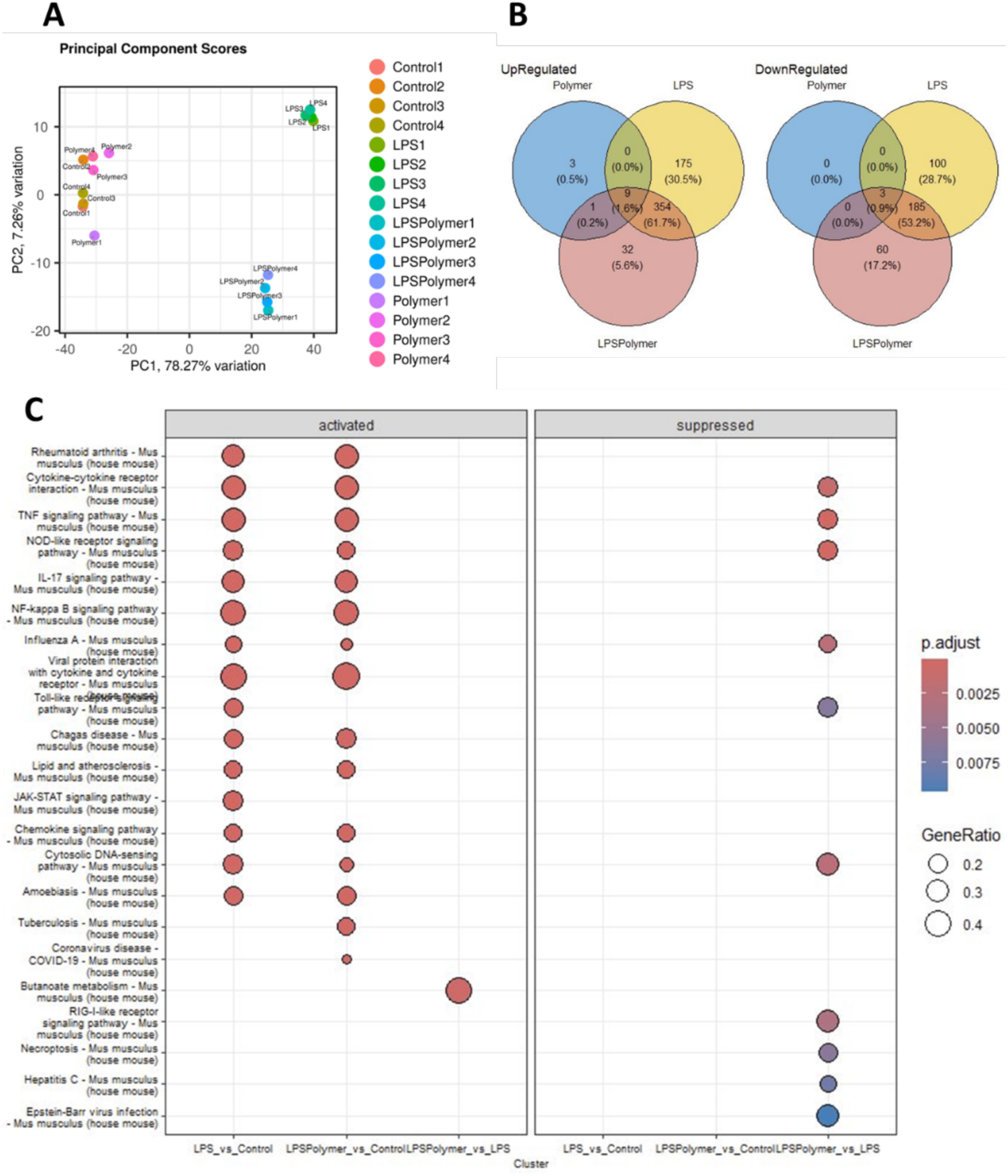
Gene expression analysis of LPS activated RAW264.7 cells after treated with copolymer 8. A) Principal component analysis (PCA) of RNA-seq. results of the control, polymer only, LPS and LPS along with polymer, B) Venn diagram of up and down regulated genes in LPS/polymer, LPS and polymer treatment in comparison to control, C) KEGG GSE analysis.

Furthermore, the polymer alone treatment did only marginally affect the gene expression profile of the RAW macrophage cell line. This signifies that polymer treatment alone does not impact transcriptomic programmes. A gene set enrichment analysis, using KEGG pathways, also uncovered the LPS treatment activated pathways that are associated with various inflammatory processes and diseases (Figure 8C). Notably, polymer treatment suppressed the activation of these pathways (P < 0.0025), particularly those related to TNF signalling, NOD-like signalling, cytokine-cytokine receptor interactions, influenza A, cytosolic DNA sensing pathway, and RIG-1 like receptor signalling pathway. Additionally, the heatmap of inflammatory genes (Figure S10) including chemokines such as Cxcl-9, 10, 11, and cytokine regulators like tnfsf10, ifnb1, ifi47, ifih1, irf1, and 7, indicated their upregulation by LPS, which was subsequently downregulated upon polymer co-treatment. Furthermore, among the top 16 altered transcription factors (TF) activities, 8 TFs were suppressed by polymer co-treatment in LPS-activated RAW264.7 macrophages. Among these 8 TFs, the key TFs involved in the LPS induced inflammation are IRF and NF κB pathway (Figure S11).

Certain key genes regulated by IRF and NF-κB are CXCL10, IRF2, IFNB1, and ISG20. These are involved in the progression various inflammatory diseases such as Alzheimer disease,^46^ inflammatory bowel diseases,^47^ and sepsis.^48^ Interestingly, these genes were found to be downregulated in the polymer treated LPS activated RAW 264.7 cells. This underscores that polymer treatment, in conjunction with LPS, exerts an immunomodulatory effect by influencing the expression of genes associated with inflammatory pathways, including the IRF and NF-κB pathways. These results point towards a potential intracellular effect of this polymer, which would indicate that the polymer does not prevent inflammation solely by binding to LPS but also by directly modulating intracellular processes.

## Conclusion

The aim of this study was to determine if we could mimic the dual effects of antimicrobial and immunomodulatory effects of CHDPs with polymers. We successfully determined the immunomodulatory properties of a library of polyacrylamides with systematically varied molecular weight, monomer distribution and composition in order to tune their hydrophobicity and cationic properties. It was found that cationic units are essential for the polymers to exert anti-inflammatory effects, inhibiting LPS-induced IL-6 secretion by human THP-1 cells by NF-kB activation in mouse RAW 267.4 cells. Furthermore, a diblock structure along with a lower molecular weight showed a good balance between significant anti-inflammatory effects and low cytotoxicity. Consequently, two lead compounds emerged, cationic diblock copolymers 3 and 8 with either hydrophobic pNIPAm (3) or hydrophilic pDMA (8) block segments, of which only the hydrophobic-cationic diblock copolymer p(NIPAm-b-AEAm)_50_ (3) exhibited significant antimicrobial activity against *P. aeruginosa*. The anti-inflammatory effects were further confirmed by differential gene expression in LPS-treated RAW 264.7 macrophages, showing that this polymer directly exerts an immunomodulatory effect on the IRF and NF-κB pathways, thereby potentially acting both as an anti-endotoxin binding to LPS as well as an intracellular modulator of inflammatory pathways. Further investigation of the mode of action is warranted to confirm these promising results, and this compound could prove effective in combating multi-resistant bacterial infections while simultaneously preventing chronic inflammation and sepsis.

## Acknowledgments

S.L. acknowledges the Commonwealth Scientific and Industrial Research Organisation (CSIRO) and the University of Warwick for the provision of a scholarship, the Polymer Characterisation Research Technology Platform at the University of Warwick for use of the SEC instruments, Azenta Life Sciences (Germany) for the gene sequencing, Dr. Ian Hands Portman and the SLS Imaging facility at the University of Warwick for access to the confocal microscope, and Cerith Harries, Caroline Stewart and the University of Warwick Media Preparation team at the University of Warwick for preparing some of the media used for this work.

## Supplementary Information

^1^H NMR spectra of all compounds synthesized in this work, SEC traces of all linear copolymers, additional information confocal microscopy images and differential gene expression analysis figures.

## Table of Contents Graphic

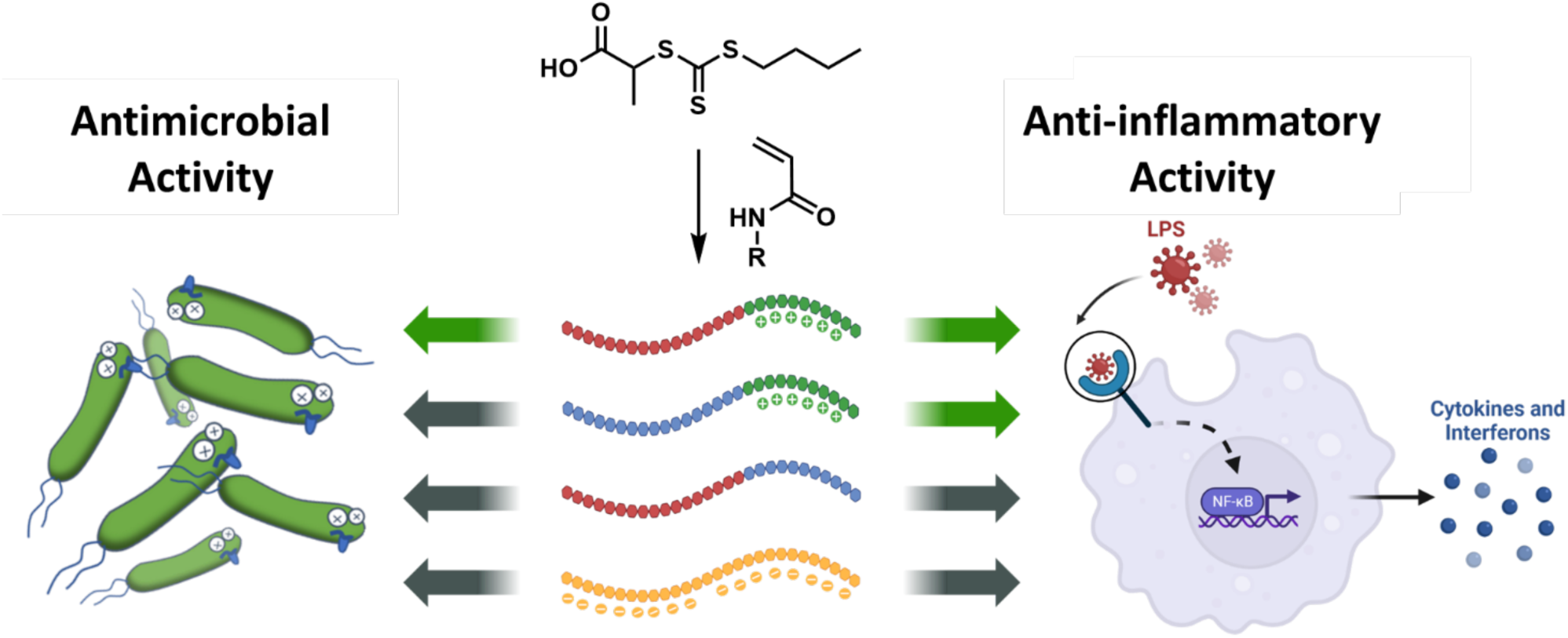

